# PARP1 inhibitors regulate PARP1 structure independent of DNA, reducing binding affinity for single strand breaks

**DOI:** 10.1101/2025.08.30.673248

**Authors:** Mark Pailing, Lotte van Beek, Taiana Maia de Oliveira, Maria M. Flocco, Bart W. Hoogenboom

## Abstract

Cancers caused by mutations to the DNA repair machinery may be treated by inhibitors that target Poly(ADP-ribose) Polymerase 1 (PARP1). PARP inhibitors are thought to cause toxicity by trapping PARP1 at single strand breaks, preventing single strand break repair, thus leading to accumulation of DNA damage and cancer cell death. Intriguingly though, different PARP inhibitors display similar cellular toxicities and catalytic inhibition despite having widely varying trapping potencies. To better understand this apparent contradiction and identify complementary mechanisms of action, we here visualize the effect of inhibitors on individual PARP1 and PARP2 molecules by atomic force microscopy (AFM). We find, surprisingly, that inhibitors cause significant PARP1 compaction and loss of molecular flexibility also in the absence of DNA. This compaction correlates with the trapping potency of the inhibitor; and could be functionally relevant by reducing the subsequent binding of pre-treated PARP1 to DNA. Such changes are less pronounced for PARP2, which shares a high sequence identity with the PARP1 catalytic domain but lacks the DNA binding domain present in PARP1. Our findings reveal an additional, DNA-independent mechanism of action for PARP inhibitors, where PARP inhibitors with strong trapping potencies target PARP1 in the absence of DNA, compact their conformation and thereby reduce its ability to bind to DNA.

## Introduction

Poly ADP-Ribose Polymerase 1 (PARP1) is part of a family of 17 proteins with various functions in the cell. PARP1 was the first member found and is primarily involved in the response to DNA damage, particularly single-strand breaks. Other roles of PARP1 include transcription regulation, chromatin remodelling, telomere maintenance and a specific cell death pathway called Parthanatos (Zandarashvili et al, 2020; Hopkins et al, 2015; Dawicki-Mckenna et al, 2016; Langelier et al, 2018). Upon binding to DNA damage, PARP1 becomes activated and produces branched poly-ADP-ribose chains on itself and target proteins, termed auto- and trans-PARylation respectively. This creates a region of high negative charge, which results in recruitment of repair proteins, release of PARP1 and repair of the DNA damage (Eustermann et al, 2015; Reber et al, 2022).

PARP1 consists of four domains further divided into sub-domains and is assumed to exist as a “beads on a string” model containing multiple folded domains joined by flexible amino acid linkers (Lilyestrom, JMB, 2010; 2018 eLife Johannes). Firstly, the N-terminal DNA binding domain (DBD) consists of 3 zinc fingers. Zinc fingers 1 and 2 bind single-strand DNA breaks (Eustermann, Mol Cell, 2015) and double-strand DNA breaks (Ali et al, Nat Struct Mol Biol, 2012) and the third atypical zinc finger 3 (Znf3) is critical for binding single-strand breaks and DNA-dependent PARP1 activation (Langelier et al, 2010). Secondly, the auto-modification domain contains the BRCA1 C-terminal domain (BRCT) where ADP-ribosylation occurs upon PARP1 activation (Dawicki-Mckenna et al, 2016; Mao et al, 2011; Altmeyer et al, 2009). The third domain is the WGR domain, named for the Tryptophan (W)-Glycine (G)-Arginine(R) rich repeat; it is critical for binding the 5’ blunt end of damaged DNA (Langelier et al, 2012; Langelier et al, 2018; Matta et al, 2020). The fourth domain is the catalytic domain (CAT), which is highly conserved among all PARPs involved in DNA damage repair (PARP1, 2 and 3). The catalytic domain is divided into the helical domain (HD) and the ADP-ribosyl transferase (ART) domains. Within the ART domain the HYE motif is conserved across the PARP family and is responsible for ADP-ribosylation catalysis. Current data suggests that when inactive, the helical domain of PARP1 sits within the catalytic domain, preventing NAD^+^ binding and locking PARP1 in an off-state. Upon binding to DNA, large scale conformation changes occur within PARP1, starting in the DNA binding domain which stacks against DNA. This causes stable cross-domain contacts to form, resulting in conformational changes across the whole protein. Critical to these is the extraction of the helical domain from the catalytic domain, opening the catalytic pocket and allowing NAD^+^ to bind and PARylation to occur (Dawicki-McKenna et al, 2016).

PARP1 inhibition was first shown in the early 2000s using non-hydrolysable NAD analogues that cannot be used for PARylation. PARP inhibitors have since been extensively studied as therapeutic agents for cancers due to BRCA1/2 mutations. PARP inhibitors cause cellular death through synthetic lethality (Hopkins et al, 2019). Synthetic lethality occurs when the loss of function of two genes causes cell death, but the loss of only one gene does not kill the cell (Lord and Ashworth, 2017). BRCA1/2-mutated cancer cells lack homologous recombination as a DNA repair pathway, which results in a stronger dependence on PARP1 to maintain genetic integrity. Hence inhibiting PARP1 in these cells causes DNA repair mechanisms to fail, resulting in genetic instability and eventually leading to gross chromosomal damage and cell death. More recently, PARP inhibitors have been trialled to treat cancers that lack other homologous recombination factors. Today, five clinical PARP1 inhibitors exist as treatments for a wide variety of cancers (Kim and Nam, 2022).

PARP inhibitors mediate their anticancer properties through a process termed PARP trapping, as follows (Hopkins et al, 2015). PARP inhibitors compete with NAD^+^ for the active site of PARP bound to single strand breaks. When the PARP catalytic site is inhibited, PARP1 cannot add Poly(ADP-ribose) chains to itself or to other downstream targets that enable DNA repair. Hence, single strand breaks go unrepaired and are converted to double strand breaks. In cells with homologous recombination defects, these cells accumulate DNA damage resulting in genetic instability, gross DNA damage and thus cell death (Hopkins et al, 2019). It was also noted that PARP complexes trapped at DNA damage sites are more cytotoxic than the naked DNA damage event. Since the PARP1 catalytic site is thought to be closely associated to the helical domain in the non-DNA bound fraction, DNA binding promotes the opening of the catalytic pocket, facilitating inhibitors binding to PARP1 (Dawicki-Mckenna et al, 2015). Recent data has classed PARP inhibitors into three groups depending on their ability to cause conformational changes in PARP1. Stronger potency trappers, such as talazoparib and olaparib, appear to be allosterically silent, whereas the weaker trappers cause allosteric changes that enhance PARP1 release from DNA (Zandarashvili et al, 2020). An overall intriguing aspect is that different PARP inhibitors have very similar toxicities and catalytic inhibition despite a wide range of trapping potency. Although PARP inhibition has been extensively studied and PARP trapping is widely accepted, it is still debated if PARP trapping is caused by catalytic inhibition, allosteric regulation or a combination of both (Zandarashvili et al, 2020; Langelier et al, 2012; Kanev et al, 2024).

This ambiguity remains in large part due to the flexible nature of PARP1, which makes it less amenable to traditional structural methods. Here we use atomic force microscopy (AFM), complemented by dynamic light scattering (DLS) and fluorescence polarisation, to examine the effects of PARP inhibitors on PARP1 and 2 in the absence of DNA; and study how binding of inhibitors to PARP1 affects subsequent binding to damaged DNA. We find that PARP inhibitors bind to PARP1 in the absence of DNA and cause major conformational changes in PARP1 independent of DNA binding. Remarkably, this binding of inhibitors to free PARP1 reduces the ability of PARP1 to subsequently bind DNA. Given that the DNA unbound fraction within the cell is abundant and participates in roles beyond DNA repair, it is possible that the clinical efficacy of the inhibitors could be partly attributed to their effects on the soluble, DNA-free PARP1 present in the cell.

## Methods

### Atomic Force Microscopy

AFM imaging was conducted at room temperature using a FastScan Bio microscope (Bruker, Santa Barbara). Images were collected using off-resonance tapping (“PeakForce Tapping” (PFT)) mode. Force curves were collected using a PeakForce Tapping amplitude of 10 nm. Images were collected at a PFT frequency of 8 kHz using FastScan-D-SS (Bruker) cantilevers (resonance frequency in liquid ∼110 kHz and nominal spring constant 0.25 N/m). Imaging speeds of 2.5-3.5 Hz (line rates) and PeakForce setpoints of 50-250 pN were used. All images were collected using 512 x 512 pixels. Initially, images were processed in Gwyddion 2.6 (Nečas and Klapetek, 2012) by mean plane subtraction and median line-by-line alignment. DNA and proteins were masked and images were processed by first levelling data to make surface facets point upward, mean plane subtraction and median line by line alignment ignoring masked features.

High-speed AFM imaging was conducted at room temperature using a NanoRacer microscope (Bruker, Berlin). Imaged were collected using the instrument’s AC fast mode. Images were collected using USC-F1.2-k0.15 cantilevers (Nanoworld; resonance frequency in liquid ∼1200 kHz and nominal spring constant 0.15 N/m). Imaging speeds of 300 lines/s and a setpoint of ∼7 nm was used, with a free amplitude of 8 nm. All images were collected using 256 x 256 pixels. Videos were processed using NanoLocz software (Heath, Micklethwaite and Storer, 2024).

For circularity analysis, AFM files were opened in NanoLocz software. Files were flattened using the “iterative fit peaks” function. The colour scale was altered to 0 nm to 2.5 nm for all frames in each video, where 0 nm corresponds to the mica surface. AFM images were then exported as .tiff and .gif format. tiff files were opened using ImageJ, pixel size and time scales were assigned using the image properties function. Thresholds were applied to mask individual PARP proteins. The analyse particles function was used; outlines of the masked particles and particle measurements were saved. Circularity for individual PARP proteins per frame was recorded over the time course. Mean circularity for was calculated for each PARP protein across the time course. Mean circularity of individual proteins was plotted against the reaction condition. Variance in circularity across the time course was calculated for individual PARP proteins using the “VAR.P” function in Excel.

### DNA

DNA containing a single strand break was PCR amplified from Lambda DNA (ThermoFisher; Cat no: SD0011) using the forward primer 5’-BiotinCGATGTGGTCTCACAGTTTGAGTTCTGGTTCTCG-3’ and the reverse primer 5’-BiotinGGAAGAGGTCTCTTAGCGGTCAGCTTTCCGTC-3’ using Phusion High-Fidelity DNA Polymerase (New England Biolabs; Cat No; M0530). The resulting product was a 496 base pair DNA with biotin modification at each end. The DNA contains a cut site for Nt.BsmAI (New England Biolabs; Cat No: R0121S) 172 base pairs from one end of the DNA. The DNA was digested using the manufacturer protocol to produce a single nick roughly 1/3 of the way along the DNA. The resulting nicked DNA was purified using a QIAquick PCR purification kit (Qiagen; Cat No: 28104) and stored at 4°C until use.

Fluorescein labelled dumbbell DNA for fluorescence polarization assays was purchased from IDT following sequences described by Eustermann et al, 2011. A 44-nucleotide construct (5’-Phosphate-CGGTCGA(fluor-dT)CGTAAGATCGACCGGCGCTGGAGCTTGCTCCAGCGC-3’) was ordered as it self-assembles into a dumbbell shape containing a single strand break. DNA was reconstituted in MilliQ water and incubated at 37°C for 1 hour to enable self-assembly (Eustermann et al, 2011).

### PARP binding molecules

β-Nicotinamide adenine dinucleotide (NAD) sodium salt (Sigma-Aldrich; Cat no: N0632) and β-Benzamide adenine dinucleotide (BAD) sodium salt (Sigma-Aldrich; Cat no: SML-3494) were reconstituted and stored at -20°C until use.

Veliparib (ApexBio; Cat no: A3002), Olaparib (ApexBio; Cat no: A4154) and Talazoparib (ApexBio; Cat no: A4153) were all purchased at 10 mM in DMSO and stored at -20°C until use.

### Proteins

Wild type PARP1 (amino acids 2-1014) or PARP2 (amino acids 2-583) was obtained from AstraZeneca UK. Human PARP1 (accession #AAH37545) with a V762A substitution and PARP2 (accession #NP_005475) were cloned into a pFastBac expression vector for baculovirus generation and contained an N-terminal 6His-6Lys-TEV or Avi-6His-TEV tag. After virus scale-up, recombinant proteins were expressed in Sf21 cells and SF900 ii SFM media for 48 hours, prior to cell harvesting at 3400 g (15 min), followed by storing the cell pellets at –80 °C.

Briefly, the recombinant proteins were purified using IMAC affinity purification and size exclusion chromatography on an AKTA system. Cells were resuspended in equilibration buffer (50 mM HEPES pH 7.5; 500 mM NaCl; 10% glycerol; 1 mM TCEP) supplemented with 20 mM imidazole; 1x EDTA-free protease inhibitor tablets (Roche) and benzonase to 5 mL per gram cells and stored at -70 °C for a freeze-thaw lysis. The lysate was clarified by centrifugation at 16000 rpm at 4 °C for 2 hours. Proteins were captured on a 5 mL FF NiNTA column (Cytiva), washed with equilibration buffer supplemented with 20 mM imidazole and eluted in equilibration buffer supplemented with 300 mM imidazole. The tags were left on and the proteins were further purified by size exclusion chromatography (S200) in 10 mM HEPES pH 7.5; 150 mM NaCl; 5% glycerol; 1 mM TCEP (PARP1) or 40 mM HEPES pH 7.5; 150 mM NaCl; 5% glycerol; 1 mM TCEP (PARP2) to an approximate purity of 95%. PARP1 and PARP2 were concentrated to 3-7 mg/mL, aliquoted, snap frozen in liquid nitrogen and stored at –70 °C. Protein QC included intact mass spectrometry to confirm protein identity and SDS PAGE analysis to confirm protein purity.

### PARP1/PARP2 compaction studies under AFM

250 nM PARP1 or PARP2 was incubated with 1 µM inhibitor in imaging buffer (12.5 mM NaCl; 12.5 mM HEPES; 1 mM Mg^2+^ pH 7.4) at room temperature for 30 minutes. Following incubation, 40 µL 10 mM Mg^2+^ in imaging buffer was added to freshly cleaved mica and 0.5 µL PARP – inhibitor mix was added. Samples were incubated for 10 minutes and then one 1 µm x 1 µm or 250 nm x 250nm square scan was taken per biological replicate. Images were processed as described above and a 3 nm mask was applied to mask individual PARP proteins. The measure individual grains function of Gwyddion was used to measure 10 PARP1 or PARP2 protein volumes using the zero-basis volume function per image.

### Dynamic Light Scattering

DLS experiments were conducted using an Anton Paar Litesizer 500 DLS system using a quartz cuvette (Anton Paar; cat no: 163390). DLS particle size analysis was conducted using back scattering (175°), automatic focus and optics. For each condition 20 runs with an exposure time of 10 s per run were conducted. PARP1 and PARP2 werediluted to 0.5 mg/ml (4.42 µM) in distilled water. Veliparib, Olaparib and Talazoparib were added to a final concentration of 1 µM and incubated at room temperature for 30 minutes prior to size measurement.

### PARP1-DNA complex studies

For DNA – first incubation studies, 250 nM PARP1 was incubated with 10 nM 496 base pair DNA containing a single strand break in imaging buffer (12.5 mM NaCl; 12.5 mM HEPES; 1 mM Mg^2+^ pH 7.4) for 30 minutes. Following this, 1 µM inhibitor was added and incubated at room temperature for a further 30 minutes. This incubation step was also conducted for no drug control conditions but without the addition of any extra compounds after the initial PARP – DNA incubation step. 6 mm mica disks were cleaved and 10 µL 1:1 PEG_113_-b-PLL_10_ : PLL_1000-2000_ (Alamanda Polymers; Cat No: 050-KC0107-107; Sigma; Cat No: P8920) was added and incubated in a humidified environment for 30 minutes (Akpinar et al, 2019). Mica disks were then washed by running 1 mL MilliQ water over each disk five times. Excess solution was removed and 40 µL of imaging buffer was added. 1.5 µL PARP-DNA-inhibitor mix was added to mica and incubated for 30 minutes. Samples were washed by running 1 ml MilliQ water at each sample disk five times, excess was removed and 40 µL fresh pH 7.4 imaging buffer was added. Samples were allowed to equilibrate prior to imaging.

For PARP – inhibitor first incubation studies, 250 nM PARP1 and 1 µM inhibitor were incubated in imaging buffer (12.5 mM NaCl; 12.5 mM HEPES; 1 mM Mg^2+^ pH 7.4) at room temperature for 30 minutes. This incubation step was also conducted for no drug control conditions, without extra compounds added to PARP proteins. Following this, 10 nM 496 base pair DNA containing a single strand break was added and incubated and room temperature for 30 minutes. 6 mm mica disks were cleaved and 10 µL 1:1 PEG_113_-b-PLL_10_ : PLL_1000-2000_ was added and incubated in a humidified environment for 30 minutes. Mica disks were then washed by running 1 mL MilliQ water over each disk five times. Excess solution was removed and 40 µL of imaging buffer was added. 1.5 µL PARP-DNA-inhibitor mix was added to mica and incubated for 30 minutes. Samples were washed by running 1 ml MilliQ water at each sample disk five times, excess was removed and 40 µL fresh pH 7.4 imaging buffer was added. Samples were allowed to equilibrate prior to imaging.

Images were processed in Gwyddion 2.6 by mean plane subtraction, median line-by-line alignment, level data by facet points, and a mask was applied to identify peaks 3nm or taller with respect to the mica surface. This identified PARP1 proteins and did not mask DNA. PARP1 was considered bound to DNA only if the protein was bound between 20% - 40% from one end of the DNA.

### Fluorescence Polarization (FP) assays

For PARP1 – DNA first experiments 250 nM PARP1 or PARP2 was incubated with a 10 nM fluorescein labelled dumbbell DNA containing a single strand break for 30 minutes at room temperature in imaging buffer (12.5 mM NaCl; 12.5 mM HEPES; 1 mM Mg^2+^ pH 7.4).

Following this, increasing concentrations of PARP inhibitors (10 pM to 300 µM) were added and incubated for 30 minutes at room temperature. Finally, 1 mM NAD^+^ was added and incubated for a further 30 minutes. 50 µL sample was added to each well in a black 96-well Nunc FluoroPlate (ThermoFisher; Cat No. 13701). Fluorescence polarization was measured using a Hidex Sense plate reader with 460 nm excitation and 520 nm emission.

For PARP1 – inhibitor first experiments, 250 nM PARP1 or PARP2 was incubated with increasing concentrations of PARP inhibitors (10 pM to 300 µM) in imaging buffer (12.5 mM NaCl; 12.5 mM HEPES; 1 mM Mg^2+^ pH 7.4) for 30 minutes at room temperature. Following this, 10 nM fluorescein labelled dumbbell DNA containing a single strand break was added and incubated for 30 minutes at room temperature. Finally, 1 mM NAD^+^ was added and incubated for 30 minutes. 50 µL sample was added to each well in a black 96-well Nunc FluoroPlate. Fluorescence polarization was measured using a Hidex Sense plate reader with 480 nm excitation and 520 nm emission.

For recovery experiments, 250 nM PARP1 was incubated with 10 µM Talazoparib for 30 minutes in imaging buffer (12.5 mM NaCl; 12.5 mM HEPES; 1 mM Mg^2+^ pH 7.4) at room temperature. Following this, 10 nM fluorescein labelled DNA containing a single strand break was added and incubated for a further 30 minutes. 1 mM NAD^+^ was added and incubated for 30 minutes. Finally, fresh untreated PARP1 (concentrations indicated in Fig. 4C) was added and incubated for 30 minutes at room temperature. 50 µL sample was added to each well in a black 96-well Nunc FluoroPlate. Fluorescence polarization was measured using a Hidex Sense plate reader with 480 nm excitation and 520 nm emission.

Dose-response curves for fluorescence polarization were fitted using a shared bottom asymptote but individual upper asymptotes.

### Statistics

Statistics and graphs were produced using Origin Pro 2023b. Student’s t-tests were used to calculate significance factors. All errors bars shown represent 1 standard deviation either side of the mean.

## Results

### PARP inhibitors bind to PARP1 and cause PARP1 compaction in the absence of DNA

Three PARP inhibitors were chosen for this study. Veliparib is known as the weakest trapper, olaparib is an intermediate trapper, and talazoparib is the strongest trapper (Zandarashvili et al, 2020). In the absence of DNA, AFM shows PARP1 as globular protrusions on the surface; and suggest a more compacted conformation following exposure to PARP inhibitors (Fig. 1A). Variable shapes are seen, in different orientations as expected for random deposition on the AFM substrate (thereby excluding AFM tip artefacts as a major factor in the overall analysis). In statistical analyses of the AFM images, PARP inhibitors (Fig. 1B) showed significant compaction of PARP1 upon incubation with inhibitors (veliparib, olaparib and talazoparib) compared to untreated PARP1. Interestingly the degree of PARP1 compaction correlates with the trapping potency of the inhibitor, as olaparib and talazoparib compacted PARP1 to a much greater degree than veliparib; the difference between higher potency trappers olaparib and talazoparib was not significant. To verify that these observations were not specific to the AFM preparation or measurement, we confirmed the same trend by DLS of PARP1 in solution, finding again that higher trapping potency inhibitors led to greater reductions in PARP1 diameter (Fig. 1D and Supplementary Table S1).

**Figure 1.**
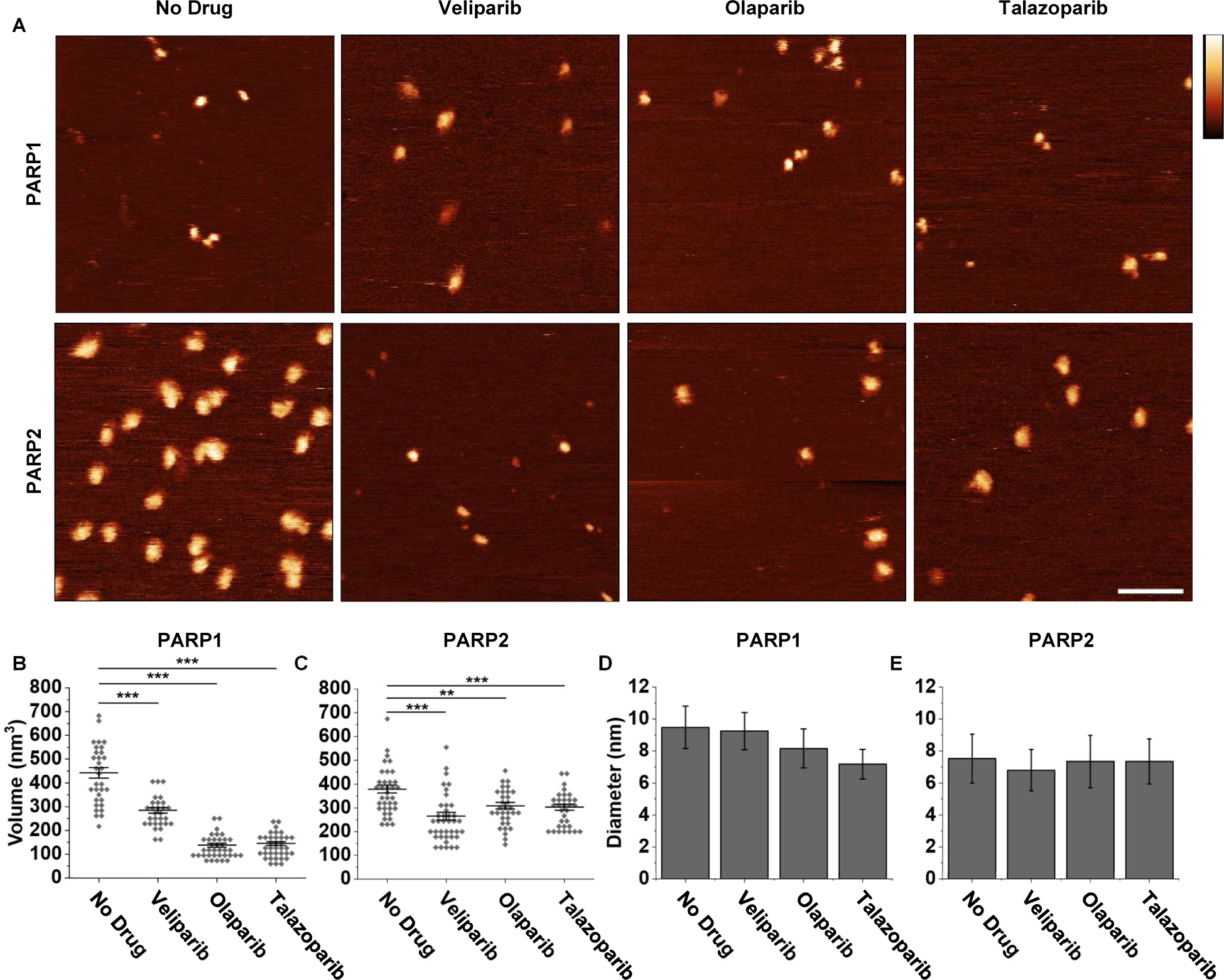
PARP inhibitors compact PARP1, and to a lesser degree PARP2, in the absence of DNA. A) AFM images of PARP1 and PARP2 adsorbed to mica following incubation with 1 µM PARP inhibitors (vertical scale = 5 nm; horizontal scale = 50 nm). B) 3D volume, as measured by AFM, of PARP1 adsorbed to mica following incubation with 1 µM PARP inhibitors (40 individual proteins spread over 4 independent AFM experiments; Biological replicates n = 4). C) As B, for PARP2. D) PARP1 diameter following incubation with 1 µM PARP inhibitors, measured using DLS (biological replicates n = 1). E) As D, for PARP2. Error bars represent 1 standard deviation from the mean. Statistical significance was quantified via t-tests: * represents p < 0.05, ** represents p <0.01, *** represents p < 0.001.

To understand which protein domains are responsible for the observed structural changes, we repeated the above experiments with PARP2. PARP2 lacks the large DNA binding domain present in PARP1 (Amé et al, 1999). The PARP2 catalytic domain has the same NAD^+^ binding motif (Karlberg et al, 2010) and is known to bind PARP inhibitors at the concentrations used here. Overall, AFM images show that preincubating PARP2 with PARP inhibitors appears to lead to some compaction of PARP2, but the differences are less pronounced (Fig. 1A). AFM data showed that veliparib, the weakest potency trapper, produced the greatest PARP2 compaction (Fig. 1C). As for PARP1, the trends observed for PARP2 by AFM were reproduced in DLS measurements (Fig. 1E). Although PARP2 has roughly half the number of amino acids of PARP1, PARP2 volume (and diameter) measurements were not half those of PARP1. This is consistent with previously published AFM measurements of PARP1 and PARP2 height (Sukhanova et al, 2016) and may be due to the inherent uncertainty in AFM volume estimates for soft and flexible samples (Giblin-Burnham et al, 2024). Overall, the difference between PARP1 and PARP2 here suggests that in PARP1, a large proportion of the compaction of the protein occurs within the DNA binding domain not found in PARP2.

### High-speed AFM highlights inhibitor-induced PARP1 conformational regulation

The observed compaction suggests that upon exposure to inhibitors, PARP1 changes from an extended, flexible and thereby presumably mobile state to a more compact, static confirmation. To test this hypothesis, we performed AFM at a higher temporal resolution (high-speed AFM, HS-AFM) on PARP1 and PARP2 with and without prior exposure to inhibitors (Fig. 2A; Supplementary Videos 1-6). To quantify the flexibility of PARP1 proteins in solution, we performed circularity analysis. Circularity is defined such that a value of 1 indicates the protein outline is identical to a circle with the same circumference as the protein outline, whereas lower values imply more irregular shapes as may be expected for more extended protein domains. Supplementary Fig. 1 shows the circularity analysis pipeline for PARP proteins. Figure 2B shows the mean circularity value for individual untreated PARP1 and 2, as well as PARP1 and 2 treated with various compounds. The variation in the circularity was calculated for individual proteins across the time course, shown in Figure 2C: It is a measure of fluctuations in the protein shape.

**Figure 2.**
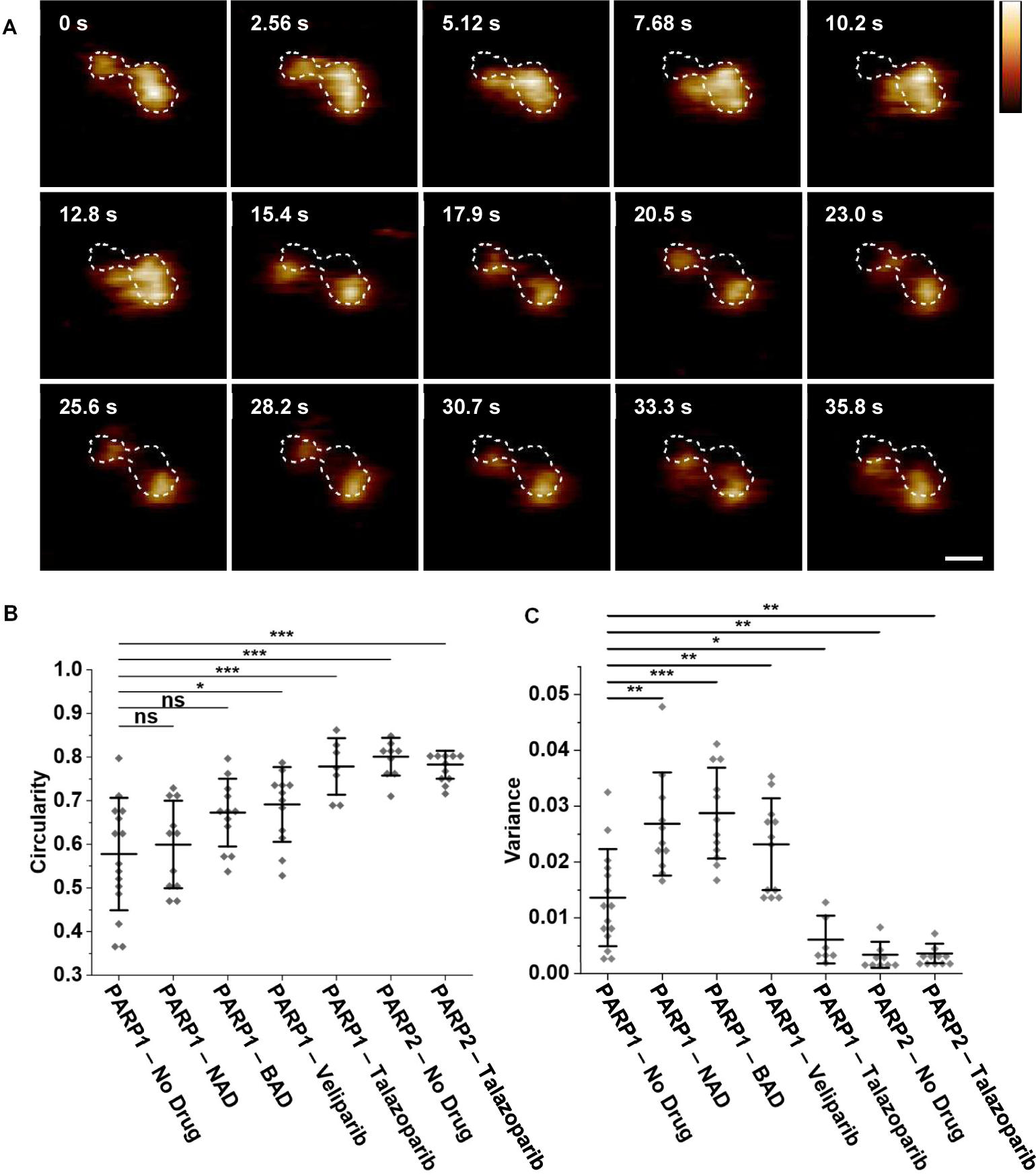
Talazoparib reduces PARP1 conformational flexibility but does not influence PARP2 flexibility. A) Extract from high-speed AFM image sequence of untreated PARP1 adsorbed to mica, illustrating PARP1 mobility in the absence of inhibitors. The outline of PARP1 from frame 1 is shown by the dotted white line (vertical scale = 2.5 nm; horizontal scale = 10 nm). B) Mean circularity of individual untreated PARP1/2 or PARP1/2 proteins treated with various compounds across a time course. Compound indicated in the figure (n = 3 biological repeats; n ≥ 7 individual proteins). C) Variance in circularity of individual PARP1/2 proteins from B across a time course (n = 3 biological repeats; n ≥ 7 individual proteins). Error bars represent 1 standard deviation from the mean. T-tests were calculated * represents P < 0.05, ** represents P <0.01, *** represents P < 0.001.

Untreated PARP1 had a large range of circularities (Fig. 2B) and a large variance in circularity (Fig. 2C) indicating that untreated PARP1 is conformationally flexible in solution. (see Supplementary Video 1). We repeatedly observed that untreated PARP1 proteins have a two lobed structure, often with a higher more defined and more stable lobe, and a smaller lobe that explores various different conformations (Fig. 2A; Supplementary Videos 1A-1D). We also observed a number of PARP1 proteins in a more extended structure (Supplementary Video 1) and in a more compacted single lobed globular structure (Supplementary Videos 1E-1F). Whilst we could roughly categorise PARP1 into these three conformation groups (single lobed, two lobed and extended), the data in Fig. 2A primarily demonstrate that the individual proteins are flexible and can swap between different conformations. For example, across the time course in Fig. 2A, the PARP1 molecule starts as a two lobed structure, by t = 25.6 seconds, it has a single lobed structure, and by t = 51.2 seconds it has gone back to the two lobed structure.

Figure 2B shows that when PARP1 is treated with NAD, BAD (a non-hydrolysable NAD analog), or veliparib the mean circularity increases compared to untreated PARP1 (Supplementary Videos 2, 3). Figure 2C shows that the variance also increased when treated with NAD, BAD and veliparib. This suggests a small degree of PARP1 compaction but also conversely an overall loosening of PARP1 stability.

High-speed AFM videos show that PARP1 treated with talazoparib occupies a static, circular structure (Supplementary Video 4). Whilst the majority of proteins had a single lobe structure, we did identify some proteins that occupied the two lobe or extended conformations. Figure 2B and 2C show that the mean circularity and variance of PARP1 are significantly reduced when treated with talazoparib. Taken together with Figure 1, this indicates that upon binding to talazoparib, PARP1 undergoes a large-scale compaction that stabilises the PARP1 conformation. Finally, the conformational change resulting in PARP1 compaction is believed to be consistent across proteins, as indicated by the reduced spread in PARP1 circularity values.

As expected, given the absence of the large DNA-binding domain (compared with PARP1), untreated PARP2 only has a single lobed structure (Supplementary Video 5) resulting in a higher mean circularity. The reduced variance in circularity also indicates that untreated PARP2 has less conformational flexibility than untreated PARP1 (Fig. 2C). Exposure to talazoparib did not lead to large changes in protein structure (Supplementary compilation video 6). PARP2 mean circularity or variance did not change greatly upon exposure to talazoparib (Fig 2B and 2C). The smaller impact on PARP2 flexibility is consistent with the smaller (compared with PARP1) changes observed in the overall size analysis (Fig. 1).

Based on the observed compaction and change in protein flexibility, and on the sequence differences between PARP1 and PARP2, we conclude that exposure to stronger trapping potency inhibitors leads to a collapse of the PARP1 DNA binding domain towards the catalytic domains, resulting in a more compact and rigid structure. Since this compaction and stabilisation of PARP1 was not seen during exposure to NAD or BAD, we conclude that binding of stronger trapping potency inhibitors induces different conformational changes in the PARP1 catalytic domain than NAD+.

### Preincubation of PARP1 with olaparib and talazoparib reduces PARP1 binding to single strand breaks

We next hypothesized that the observed loss in protein flexibility would be accompanied by a loss in protein function. Specifically, we suggest that a collapse of the DNA binding domain towards the catalytic domain and a reduction in flexibility may result in less efficient binding to single strand breaks in DNA. To test this, we incubated PARP1 with inhibitors before adding DNA with a single strand break (PARP – inhibitor first incubation). The results of these incubations were compared with those of a preparation in which PARP1 and DNA with a single strand break were incubated first before adding the inhibitors (PARP – DNA first incubation).

First, we analyzed PARP1-DNA binding at a single-molecule level by counting how many DNA molecules showed a PARP1 bound at the single-strand break located within 25% and 45% of a DNA end for the construct used here (Fig. 3A; Supplementary Fig. 4). Notably, for inhibitors causing the strongest compaction, olaparib and talazoparib, (see Fig. 1), exposing PARP1 to these inhibitors before DNA resulted in a 4 and 6-fold reduction, respectively, in the subsequent binding to DNA, compared with protocols in which PARP1 was allowed to bind to DNA before exposure to inhibitors (Fig. 3B). By contrast, for veliparib, the change in binding between the two protocols was not significant, suggesting that any changes in PARP1 due to veliparib are too minor to significantly influence PARP1 ability to bind to damaged DNA. During untreated controls there was an increase in PARP1 binding during PARP – inhibitor first incubation (Fig. 3B). Since DNA adsorption to mica is driven by electrostatic interactions DNA adsorption is often not uniform across a whole mica disc (Akpinar et al, 2019). To account for this, multiple areas of each replicate were analyzed, however this may result in the observed difference between the two protocols.

**Figure 3.**
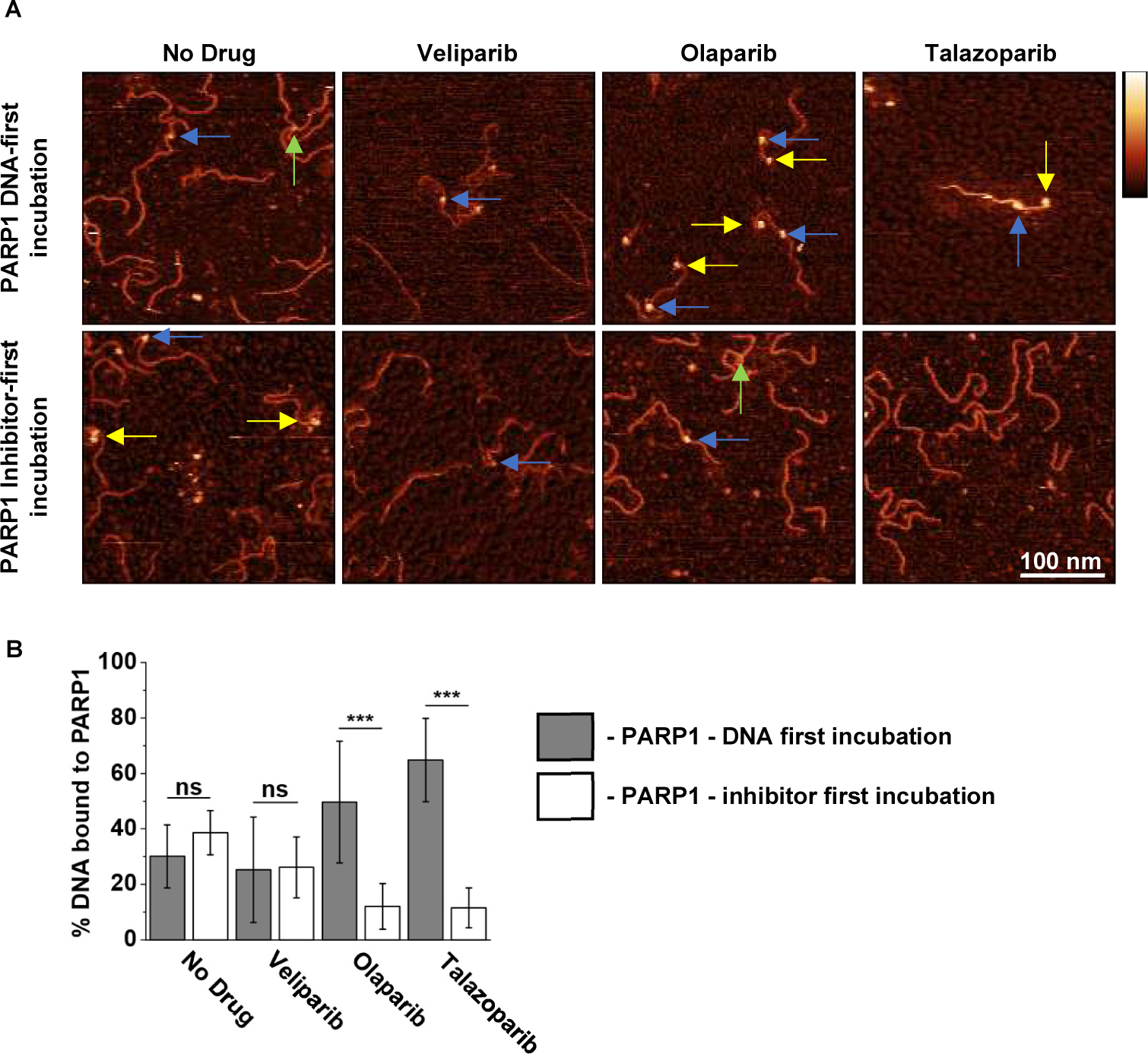
Incubating PARP1 with olaparib and talazoparib reduces subsequent PARP1-DNA binding. A) AFM images of PARP1-DNA binding, where blue arrows indicate PARP1 bound to DNA within 25-45% of a free DNA end, noting that single strand break site is at 172 bp, of the total 496 bp of the DNA, hence at about 1/3 from a DNA free end. Blue arrows indicate PARP1 presumed bound to single strand break. Yellow arrows indicate PARP1 bound to a free DNA end. Green arrows indicate sites where two DNA strands have crossed over each other giving the impression of a tall feature along the DNA backbone. In the top row (“PARP1 – DNA first”), PARP1 and DNA have been incubated prior to inhibitor treatment. In the bottom row (“PARP1 – inhibitor first”), PARP1 has been treated with inhibitors and next incubated with DNA. (Vertical scale = 5 nm; horizontal scale = 50 nm). B) Percentage of DNA molecules with PARP1 located within 25% and 45% of a free DNA end (2 AFM images for each of *n* = 4 biological preparations). Error bars represent 1 standard deviation from the mean. Statistical significance was quantified via t-tests were calculated * represents p < 0.05, ** represents p <0.01, *** represents p < 0.001.

### Inhibitor-induced reduction of PARP1-DNA binding is confirmed in bulk assays

The inhibitor-induced reduction in PARP1-DNA binding was confirmed by bulk fluorescence polarisation assays, in which the fluorescence polarisation signal is a measure of PARP1-DNA binding and in which there is no need to adsorb the samples to an AFM substrate (Moerke, 2009). In these assays, the PARP – DNA first and PARP – inhibitor first incubation protocols were adapted from the above AFM experiments to include NAD^+^ (Supplementary Fig. 2), where the presence of NAD^+^ ensured that PARP would be released from the DNA unless it was trapped on the DNA by inhibitors. Based on the AFM experiments above, we predicted that prior exposure to olaparib and talazoparib would reduce the maximum fluorescence polarization signal as measured at high PARP concentrations, since the inhibitor-bound PARP1 would have reduced affinity for the DNA. Fluorescence polarisation data was collected for PARP1 – DNA first and PARPi – inhibitor first protocols and is shown in Figure 2B. In brief, the PARP1 – inhibitor first incubation was predicted to lead to a lower fluorescence polarization signal than the PARP1 – DNA first incubation. In line with this prediction and with the AFM experiments, the maximum fluorescence polarization signal for the PARP1 – inhibitor first incubation was only marginally smaller than for the PARP1 – DNA first incubation in the case of veliparib, and a much larger and statistically significant difference was observed for olaparib and talazoparib (Supplementary Fig. 5).

Similar changes were found for the EC_50_ values (Supplementary Table 2), which define the concentrations of inhibitors needed to increase the fluorescence polarisation signal to 50% of the maximum polarisation signal. This is assumed to represent the concentration at with there is a 50% increase in the amount of PARP1 trapped to DNA compared to the lowest concentration. Here exposing PARP1 to an inhibitor prior to DNA (PARP – inhibitor first incubation) was predicted to shift the EC_50_ to higher values compared to PARP – DNA first incubation results, since the inhibitor-induced PARP1 compaction reduces binding of PARP1 to DNA (Fig. 3). Whereas this shift appeared to be insignificant for veliparib, it was more pronounced for olaparib (116 nM to 127 nM) and talazoparib (83 nM to 317 nM). As for the AFM experiments, PARP2 served as a negative control, in which the difference between PARP2 – inhibitor first and PARP2 – DNA first incubations appeared insignificant.

Finally, to confirm that the maximum fluorescence polarisation signal in these experiments was limited by the amount of PARP molecules effective in binding to DNA, recovery experiments were carried out (Fig. 4C). In these experiments, the PARP1 – inhibitor first protocol (with a PARP1 concentration of 250 nM) was followed but with the addition of fresh, untreated PARP1 (termed “Additional PARP1” in Fig. 4C) after completing all the original preparation steps (see Supplementary Fig. 3 for schematic). Fig. 4C shows the fluorescence polarisation signal for PARP1 treated with a high (10 µM) talazoparib concentration, for PARP1 – DNA first incubation, PARP1-inhibitor first incubation and recovery protocols including the addition of different concentrations of fresh PARP1. As expected, the addition of fresh 250 nM PARP1 leads to the recovery of the fluorescence polarisation signal to slightly above the result for the PARP1 – DNA first incubation. Higher concentrations of additional fresh PARP1 resulted in even more PARP1 binding to DNA. These results confirm that at high inhibitor concentrations, the availability of native PARP1 limits the DNA-PARP1 binding in these experiments.

**Figure 4.**
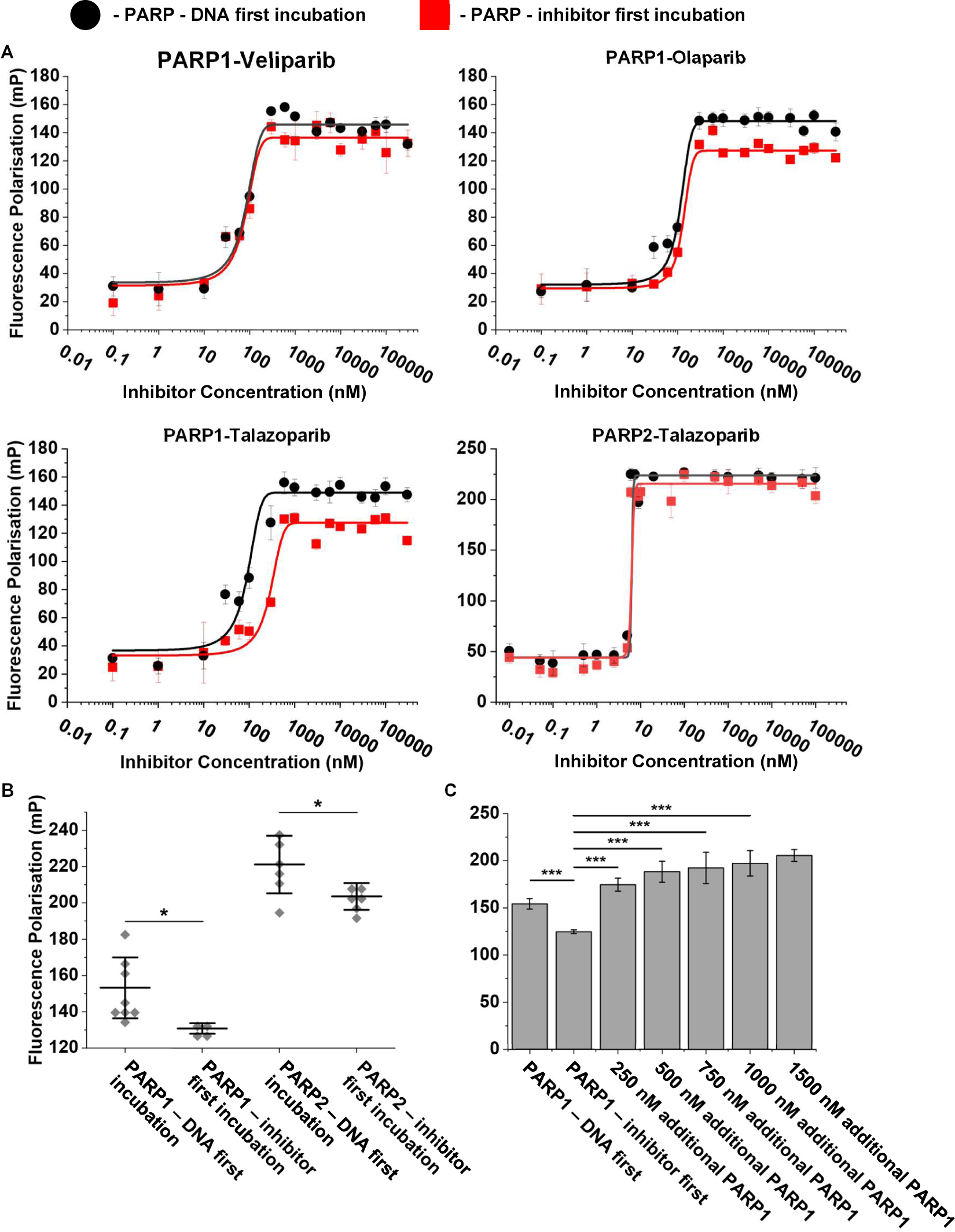
Fluorescence polarization confirms the inhibitory effect of potent PARP trappers on PARP1 binding to DNA. A) Fluorescence polarization of PARP1/2 during PARP – DNA first and PARP - inhibitor first incubation with veliparib, olaparib and talazoparib (technical replicates n ≥ 4). B) Fluorescence polarization of PARP1 and PARP2 during PARP – DNA first and PARP – inhibitor first incubation with 100 µM talazoparib and DNA containing a single strand break (n ≥ 4). C) Fluorescence polarization in binding recovery experiments, which demonstrate how the maximum PARP1-DNA binding signal recovers and further increases upon the addition of fresh PARP1 (without prior inhibitor treatment) following initial exposure to 250 nM PARP that was pretreated with 10 µM talazoparib (PARP – inhibitor first protocol) (technical replicates n ≥ 4). Error bars represent 1 standard deviation from the mean. Statistical significance was quantified via t-tests were calculated * represents p < 0.05, ** represents p <0.01, *** represents p < 0.001.

**Figure 5.**
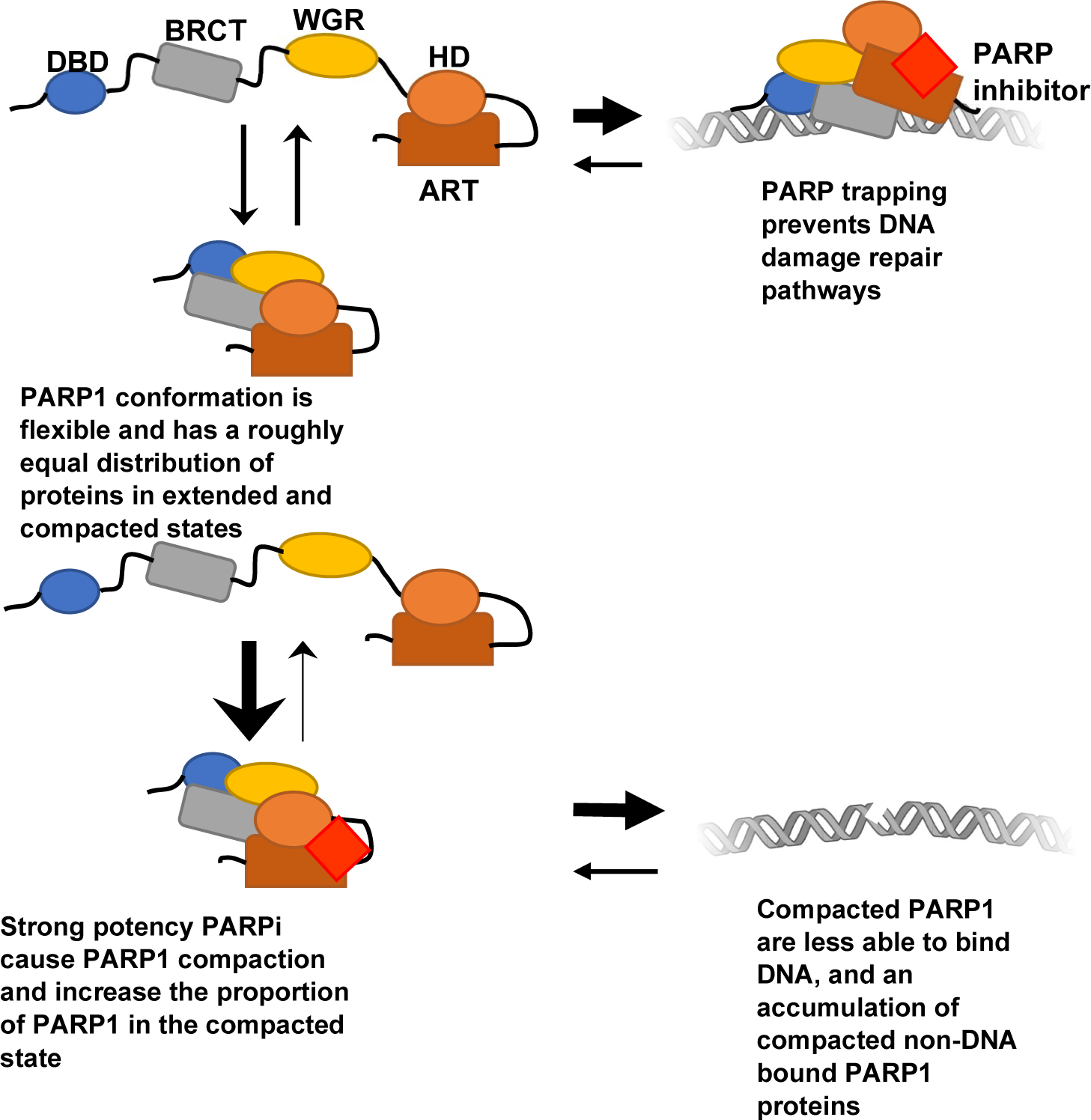
Schematic for inhibitor induced PARP1 compaction with strong potency trappers. Taken together, our observations imply an additional mechanism of action, particularly for strong trapping potency PARP inhibitors, by which inhibitors not only cause DNA-bound PARP to be trapped to the DNA, preventing further DNA repair, but also bind to free PARP to cause conformational changes that reduce the ability of PARP to bind to single strand breaks (Fig.5). This mechanism of action may co-exist with the generally assumed trapping of PARP to DNA in the presence of inhibitors and thereby shed a new light onto the complex relation between molecular trapping potency and efficacy.

## Discussion

Through the use of high-speed AFM, we present single molecule imaging of full length PARP1, providing direct evidence that PARP1 is inherently flexible. We show that PARP1 conformations are heterogenous and dynamic, whereas PARP2 adopts a more globular, uniform structure (Fig. 2). Our data suggests that the large DNA binding domain and BRCT domain present in PARP1 (but not PARP2) are the dynamic regions reported in our data. The inherent conformational flexibility may be necessary to bind to numerous types of DNA damage (Sukhanova et al, 2016; Langelier et al, 2012); this flexibility also explains why traditional structural studies (more effective for stable molecular conformations) have failed to capture the full-length structure of PARP1 (Rouleau-Turcotte et al, 2022).

Building on these observations, we have identified inhibitor-induced conformation changes of PARP1 and PARP2. Whilst previous work explores reverse allostery in the context of PARP1 retention on DNA damage sites, we explored the effects of reverse allostery in the absence of DNA. Veliparib was previously shown to destabilize the helical domain of PARP1 whilst not impacting interdomain contacts (Zandarashvili et al, 2020). Our data provides evidence that similar changes occur when veliparib binds to PARP1 in the absence of DNA.

Larger PARP inhibitors, such as talazoparib and olaparib, are known to have a higher trapping potency (Murai et al, 2012) and induce mild pro-retention reverse allosteric changes (Zandarashvili et al, 2020). These changes arise from destabilizing different regions of the helical domain compared to veliparib. Olaparib and talazoparib also promote the formation of interdomain contacts stabilizing PARP1 on DNA (Zandarashvili et al, 2020). Importantly, we have here presented evidence for reverse allosteric regulation due to PARP inhibitors binding to PARP1 in the absence of DNA. Previous studies had indicated that upon binding to DNA, interdomain contacts form between the DNA binding domain and the catalytic domain, resulting in destabilization of the helical domain and as such its removal from the catalytic pocket, thus facilitating NAD+ or inhibitor binding (Murai et al, 2012; Zandarashvili et al, 2020). Through high-speed AFM imaging, we have observed the effects of PARP inhibitors on PARP1 and 2. Inhibitors induce allosteric changes in PARP1 in the absence of DNA, causing compaction of PARP1. This correlated with reduced DNA binding of the inhibitor treated PARP1.

Since conformational flexibility is critical for binding different forms of DNA damage (Sukhanova et al, 2016; Langelier et al, 2012), reverse allosteric regulation of PARP1 accounts for the loss of PARP1 DNA binding caused by stronger potency trappers. PARP inhibitors are known to alter protein expression and reduce inflammation in a number of diseases both cancerous and non-cancerous (Teyssonneau et al, 2021; Giansanti et al, 2010). The observed inhibitor induced loss of PARP1 – DNA binding warrants further investigation as to how this population of DNA-binding deficient PARP1 proteins impacts the non-DNA repair PARP1 roles in cells.

## Acknowledgements

This work was funded by the UK Biotechnology and Biological Sciences Research Council (BB/W510026/1 to M.P. and B.W.H.; BB/W019345/1 to B.W.H.) and the UK Engineering and Physical Sciences Research Council (EP/M028100/1 to B.W.H.). The authors acknowledge Nick Bell for discussions.

## Author contributions

M.P. performed experiments and analysed the data. L.v.B., M.P., T.M.O, M.M.F., and B.W.H. designed the study and provided input on experimental analysis and interpretation. M.P. and B.W.H. wrote the initial draft of the manuscript. All authors reviewed and commented on the manuscript

## Supplementary information

**Supplementary Figure 1.**
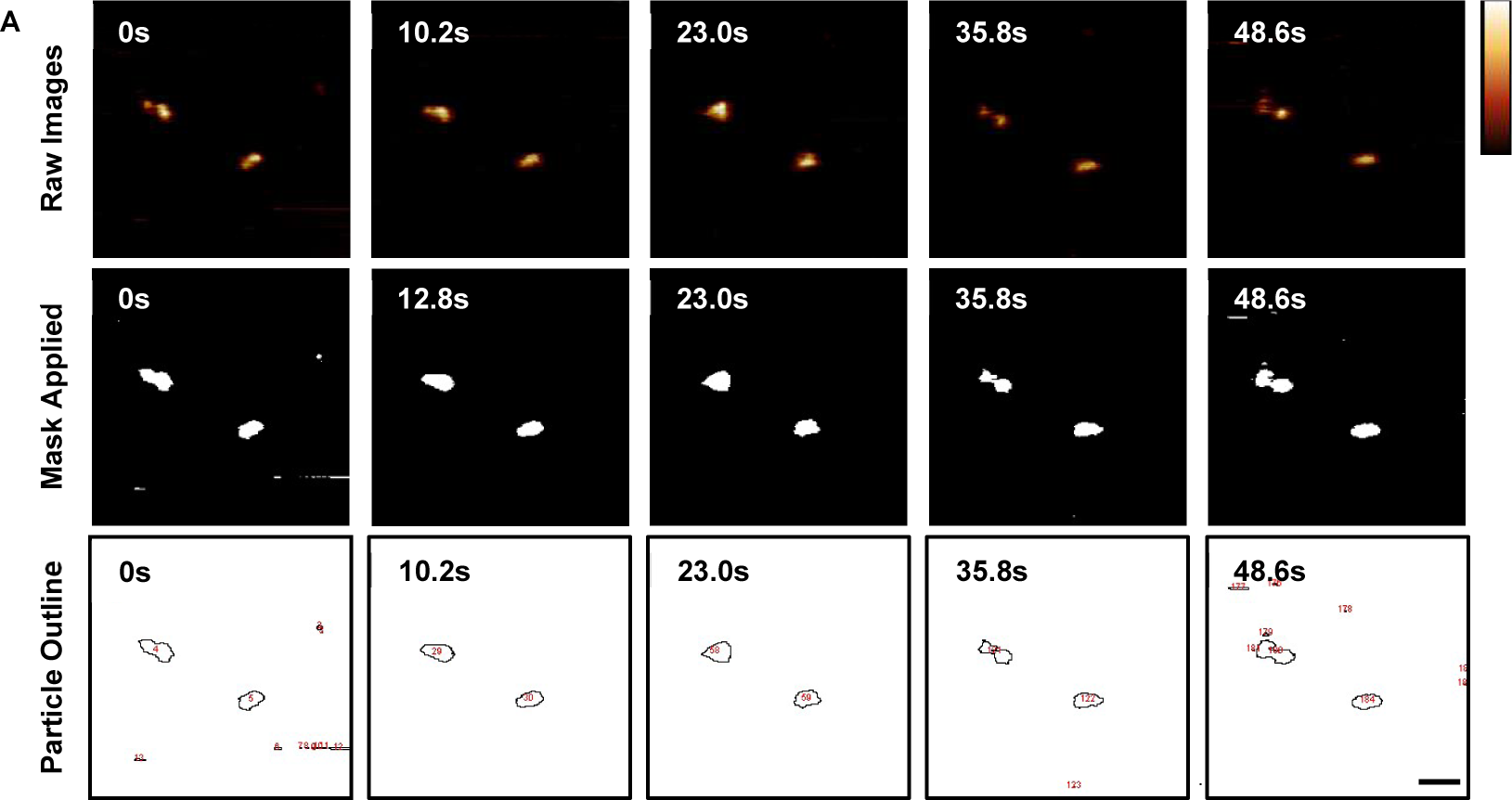
Circularity analysis pipeline of untreated PARP1 showing the raw images, protein masking and protein outlines.

**Supplementary Figure 2.**
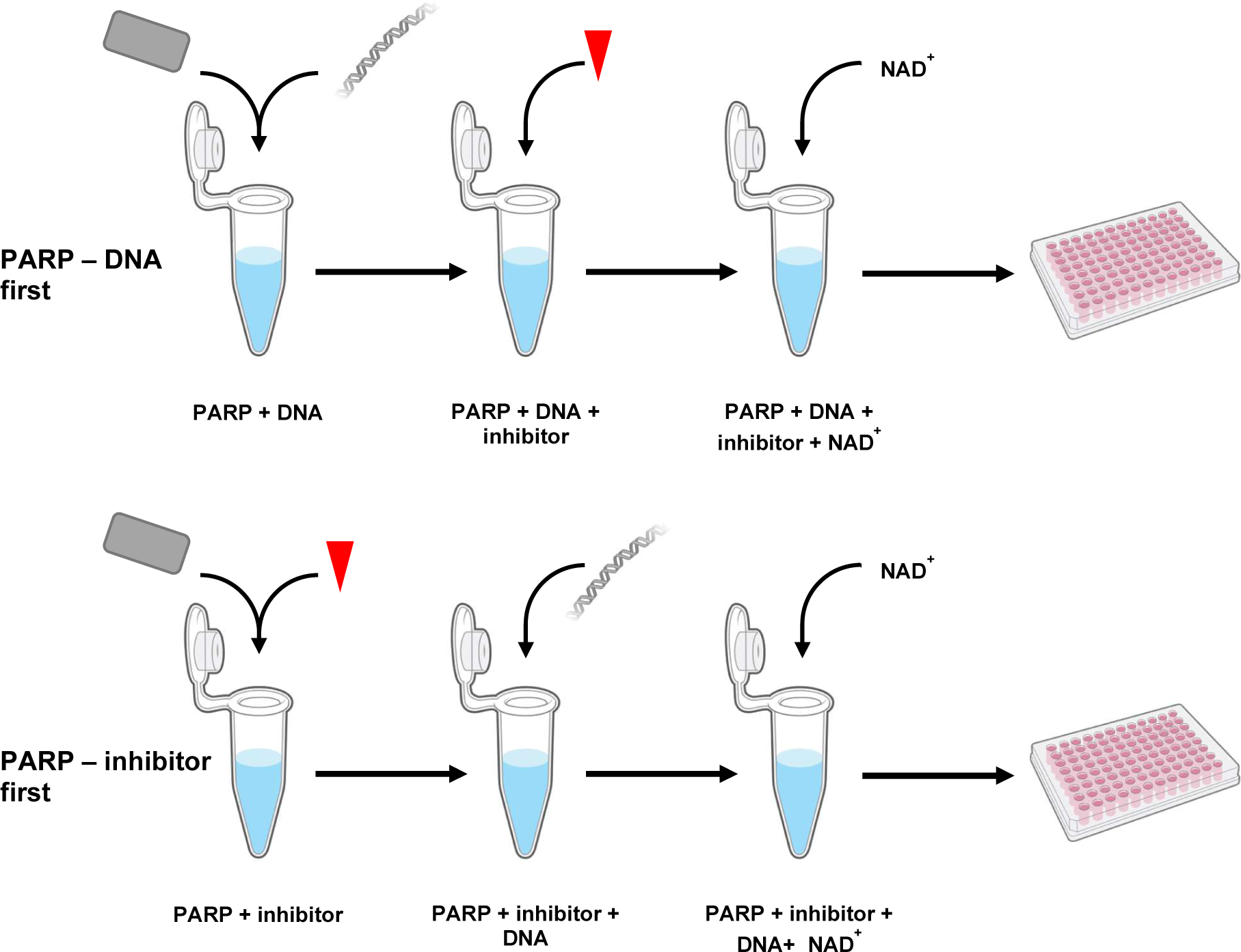
PARP – DNA first and PARP – inhibitor first protocols used for fluorescence polarisation experiments.

**Supplementary Figure 3.**
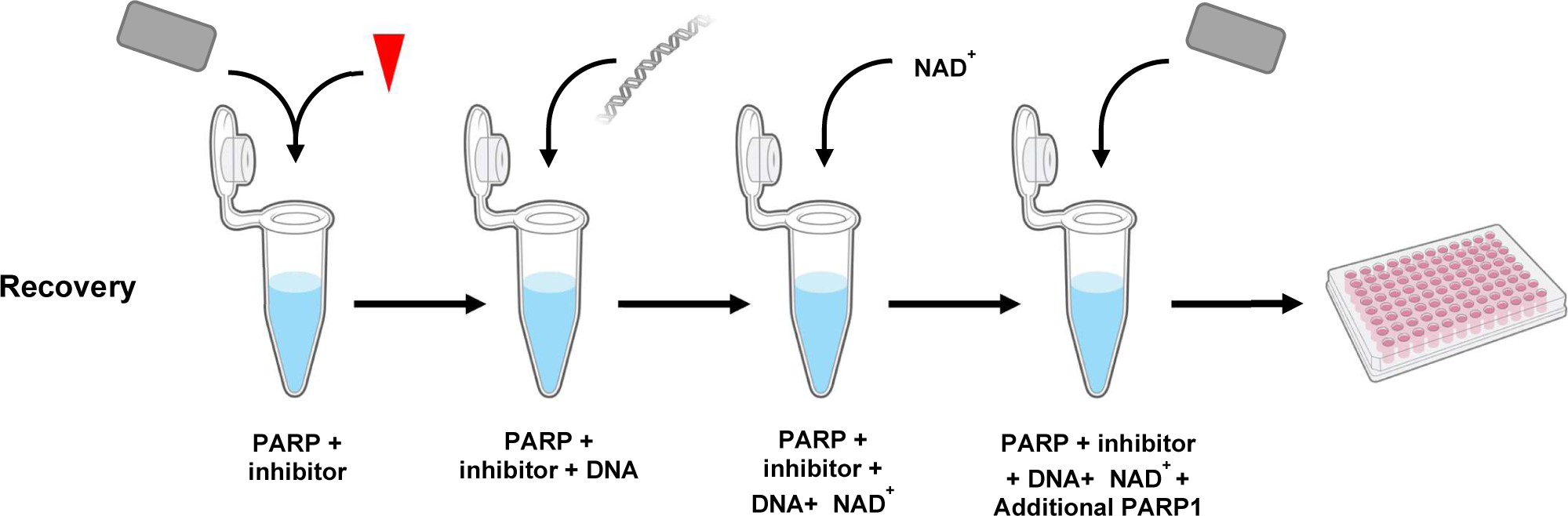
Recovery protocol used for fluorescence polarisation experiments.

**Supplementary Figure 4.**
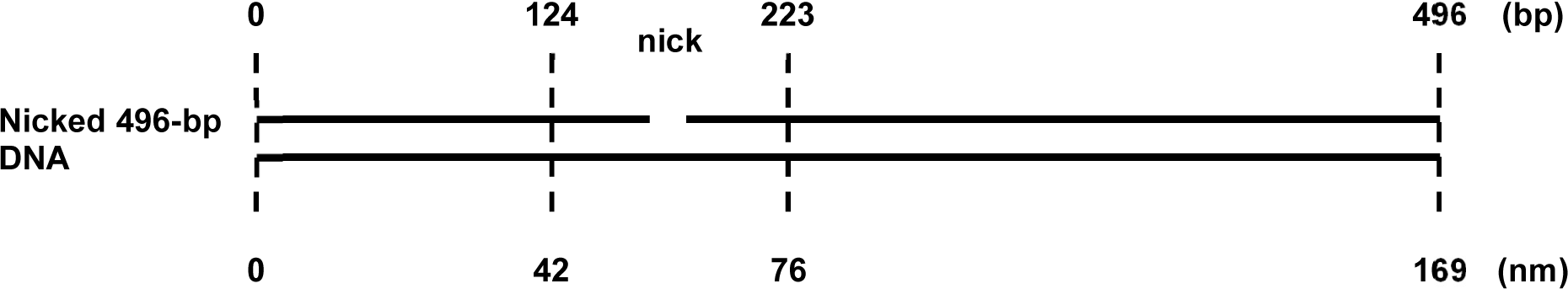
496-base pair DNA containing a single strand break. 496-base pair DNA construct containing a single strand break. PARP1 was considered bound to DNA if it was located with 25% and 45% of one DNA end. The length along the DNA of this area is indicated in nanometer.

**Supplementary figure 5.**
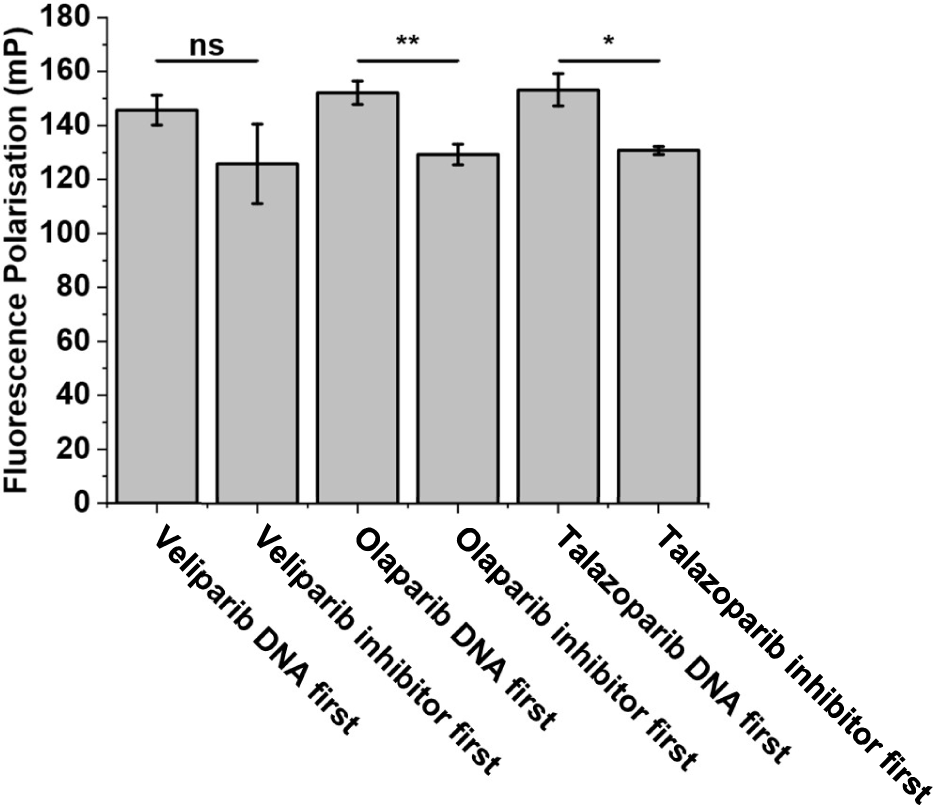
Fluorescence polarization confirms incubating PARP1 with potent PARP trappers prior to DNA reduces PARP1 binding to DNA. A) fluorescence polarisation for PARP1 incubated with 100 µM Veliparib, Olaparib or talazoparib during DNA first and inhibitor first protocols (n ≥ 4). Error bars represent 1 standard deviation from the mean. T-tests were calculated * represents P < 0.05, ** represents P <0.01, *** represents P < 0.001.

**Supplementary Table 1.** Experimentally determined PARP1 and PARP2 diameters with various inhibitors. Peak volume analysis of 4.42 µM PARP1 and PARP2 diameter treated with 1 µM inhibitors. Peak volume analysis was reported due to the complexity of protein interactions as described by Vasil’eva et al (Vasil’eva et al, 2019).

|  | Peak<br>Volume 1<br>(nm ± SD) | Area<br>Volume<br>1 (%) | Peak<br>Volume 2<br>(nm ± SD) | Area<br>Volume<br>2 (%) | Peak Volume<br>3 (nm ± SD) | Area<br>Volume 3<br>(%) |
| --- | --- | --- | --- | --- | --- | --- |
| **PARP1 No**<br>**Drug** | 9.48 ± 1.33 | 95.69 | 1072.38 ±<br>349.05 | 4.31 | N/A | N/A |
| **PARP1 +**<br>**Veliparib** | 9.25 ± 1.16 | 93.61 | 1049.32 ±<br>210.44 | 3.94 | 13724.53 ±<br>3146.77 | 2.45 |
| **PARP1 +**<br>**Olaparib** | 8.16 ± 1.23 | 94.92 | 1154.88 ±<br>387.30 | 3.25% | 6599.86 ±<br>178.08 | 3.25 |
| **PARP1 +**<br>**Talazoparib** | 7.18 ± 0.92 | 97.11 | 1228.87 ±<br>363.49 | 2.89 | N/A | N/A |
| **PARP2 +**<br>**No Drug** | 7.53 ± 1.53 | 100 | N/A | N/A | N/A | N/A |
| **PARP2 +**<br>**Veliparib** | 6.80 ± 1.29 | 100 | N/A | N/A | N/A | N/A |
| **PARP2 +**<br>**Olaparib** | 7.34 ± 1.64 | 100 | N/A | N/A | N/A | N/A |
| **PARP2 +**<br>**Talazoparib** | 7.35 ± 1.41 | 100 | N/A | N/A | N/A | N/A |

**Supplementary Table 2.** EC_50_ values for PARP1-DNA and PARP1-inhibitor first fluorescence polarisation experiments.

| Condition | EC50 (nM) |
| --- | --- |
| PARP1-Veliparib DNA First | 80 |
| PARP1-Veliparib inhibitor First | 69 |
| PARP1-Olaparib DNA First | 116 |
| PARP1-Olaparib inhibitor First | 127 |
| PARP1-Talazoparib DNA First | 83 |
| PARP1-Talazoparib inhibitor First | 317 |
| PARP2-Talazoparib DNA First | 5 |
| PARP2-Talazoparib inhibitor First | 5 |

